# A pyramidal silicon nanopore for single misfolded Tau protein characterisation

**DOI:** 10.64898/2026.08.04.742667

**Authors:** Hoi Lam Cheung, Dehua Hu, Jianxin Yang, Ho Pui Ho

**Author notes:** **Corresponding Author: Ho Pui Ho** - Ho Sin-Hang Engineering Building, The Chinese University of Hong Kong, Shatin, NT, HK.

## Abstract

Solid-state nanopores offer a versatile platform for single-molecule sensing owing to their mechanical robustness, tuneable geometry, and compatibility with scalable fabrication. Here, we present a pyramidal silicon nanopore with a 40 nm sensing aperture for label-free characterisation of protein molecules by resistive pulse sensing. The nanopore operates stably over transmembrane voltages ranging from −2 to +2 V and across a broad range of electrolyte concentrations, enabling analysis under diverse experimental conditions. As molecules traverse the confined sensing region, transient ionic current modulations are generated that reflect their excluded volume and molecular geometry. Using this approach, we characterise unlabelled Tau species spanning monomeric proteins, intermediate aggregates, and mature fibrillar assemblies. Analysis of the resulting current signatures, together with simplified geometrical models, enables reconstruction of molecular dimensions and discrimination of distinct Tau populations based on their electrical fingerprints. These results demonstrate that pyramidal silicon nanopores provide a sensitive and scalable platform for label-free monitoring of structurally heterogeneous protein aggregation and establish a framework for investigating protein aggregation using solid-state nanopore sensing.

## INTRODUCTION

Nanopore sensing has emerged as a powerful platform for single-molecule analysis, enabling label-free detection of biomacromolecules through electrical readout. Among the available nanopore platforms, biological nanopores, typically derived from membrane proteins such as α-haemolysin or MspA [1-5], provide atomically precise and highly uniform pore structure. This structural uniformity results in low device-to-device variability and well-defined transport properties, making them powerful platforms for high-resolution molecular sensing. Biological nanopores have been extensively applied in single-molecule DNA sequencing and protein detection due to their excellent signal reproducibility and intrinsic molecular selectivity. However, biological nanopores also present inherent limitations that restrict their broader applicability. Their fixed pore size, limited mechanical stability, and sensitivity to environmental conditions constrain their use for large or structurally heterogeneous biomolecules[6]. Furthermore, because the pore dimensions of biological nanopores are intrinsically defined by their protein architecture, they cannot be readily tailored to accommodate different analytes or sensing requirements. As a result, optimisation of nanopore dimensions for structurally diverse biomolecular systems remains challenging. These limitations highlight the importance of pore size as a critical parameter in nanopore sensing. Accurate discrimination of biomolecular structures often requires matching the sensing aperture to the characteristic dimensions of the target analyte in order to maximise sensitivity and information content.

To overcome these limitations, solid-state nanopores have emerged as a versatile alternative owing to their tuneable geometry, mechanical robustness and compatibility with scalable semiconducting fabrication. These systems typically consist of a nanometre-scale aperture fabricated in thin insulating membranes such as silicon nitride, silicon dioxide, or related materials[7]. When immersed in electrolyte and biased with an external voltage, ionic current flows through the nanopore, which is partially blocked when a molecule translocates through the sensing region. This transient blockade generates a characteristic resistive pulse signal, providing a physical “fingerprint” that encodes information about molecular size, shape, and conformational state[8-10]. The ability to engineer pore geometry, surface chemistry, and device architecture makes solid-state nanopores highly attractive for a broad range of applications, including nucleic acid sequencing[11-14], protein analysis[15-22], and nanoscale molecular biosensing[23-25]. In particular, their compatibility with semiconductor fabrication technologies enables scalable and robust device integration, offering significant advantages for long-term operation under diverse experimental conditions. Recent advances have enabled fabrication of pores spanning from a few nanometres to tens of nanometres[22, 26, 27], including nanopores comparable to or exceeding the dimensions of many biological assemblies[28]. Such controllability provides opportunities to optimise sensing performance across a broad range of molecular sizes and morphologies.

Protein aggregation of Tau is of particular interest in this context, as it is a key pathological hallmark of neurodegenerative disorders, such as Alzheimer’s disease and related tauopathies[29-32]. Tau proteins undergo progressive misfolding and assembly into oligomeric and fibrillar species with distinct structural and toxic properties[33-36]. Importantly, intermediate aggregation species are believed to contribute significantly to disease progression, yet remain difficult to characterise using conventional ensemble-averaged techniques because of their transient and heterogeneous nature. Recent studies have demonstrated the feasibility of applying nanopore sensing to monitor Tau aggregation and distinguish distinct aggregate populations[37]. However, quantitative structural characterisation of Tau assemblies remains challenging owing to the broad size distribution and morphological heterogeneity of the aggregates. In particular, nanopore dimensions optimised for small oligomeric species may become less suitable for analysing larger fibrillar assemblies, limiting comprehensive investigation across multiple aggregation states. In this study, we develop a pyramidal silicon nanopore platform for investigating Tau aggregation at the single-molecule level using resistive pulse sensing. By combining size-tuneable solid-state nanopores with simplified geometrical interpretation of signal features, our approach enables comparative analysis of morphologically distinct Tau assemblies and estimation of their structural dimensions. The platform provides a label-free and scalable strategy for probing aggregation heterogeneity and demonstrates the potential of engineered solid-state nanopores for analysing structurally diverse protein systems relevant to neurodegenerative disease.

## MATERIALS AND METHODS

### Preparation of Monomeric and Aggregated Tau Samples

Heparin-induced Tau aggregation samples were prepared using the protocol described by Crespo et al. [38] with slight modifications. Reaction buffer was prepared by dissolving 0.01% (v/v) Tween 80 (Sigma-Aldrich, St. Louis, MO, USA) and 0.5 mM tris(2-carboxyethyl)phosphine (TCEP; Invitrogen, Carlsbad, CA, USA) in phosphate-buffered saline (PBS, pH 7.4; Gibco, Grand Island, NY, USA), followed by adjustment of the final pH to 6.7. The solution was thoroughly mixed, filtered through a sterile 0.2 µm polyethersulfone (PES) membrane filter, and stored at −80 °C until use. A 50 µM heparin stock solution (Sigma-Aldrich, St. Louis, MO, USA) was prepared in reaction buffer and subsequently diluted to a working concentration of 2.5 µM [39].

Recombinant human Tau-441 protein [huTau441, molecular weight (MW) = 67 kDa; Novus Biologicals™, Minneapolis, MN, USA] was equilibrated to room temperature prior to use and diluted with reaction buffer. Aggregated Tau samples were prepared by mixing huTau441 with heparin in a total reaction volume of 100 µL, whereas monomeric control samples were prepared by replacing heparin with an equivalent volume of reaction buffer. The resulting mixtures were incubated in a dry bath at 23 °C before subsequent characterisation.

### Transmission Electron Microscopy Characterisation

Tau samples collected after 0, 24, and 48 h of incubation were characterised by transmission electron microscopy (TEM) to examine the morphology of Tau species formed during aggregation. A 10 µL aliquot of each sample was deposited onto a carbon film-coated 300-mesh copper grid and allowed to adsorb for 2 min before excess solution was removed using filter paper. The grids were subsequently rinsed with 10 µL of deionised water and negatively stained with 10 µL of 2% (w/v) sodium molybdate solution. After drying at room temperature for 24 h, TEM images were acquired using a JEM-2010 transmission electron microscope (JEOL Ltd., Tokyo, Japan) operated at an accelerating voltage of 200 kV. We used ImageJ[40] for TEM image analysis.

### Fabrication of Pyramidal Silicon Nanopores

Micro-pyramidal silicon nanopores used in this study were fabricated based on the closed-loop photovoltaic electrochemical etch-stop strategy previously reported by Yang et al. [26]. The fabrication process employed photoinhibition-assisted wet chemical etching in potassium hydroxide (KOH) solution with real-time electro-optical feedback, enabling precise control over nanopore formation within a 21.25 µm-thick silicon substrate. This approach allows the reproducible fabrication of pyramidal nanopores with tuneable pore dimensions spanning from sub-10 nm to tens of nanometres.

Compared with conventional nanopore fabrication techniques, the nanopore platform provides a robust solid-state architecture and a broad accessible pore-size range, making it suitable for the analysis of biomolecules with diverse dimensions. Detailed fabrication procedures and device characterisation have been reported previously by Yang et al. [26].

### Nanopore Experimental Setup

For nanopore measurements, the fabricated chip was mounted onto a custom polymethyl methacrylate (PMMA) microfluidic chamber, separating the system into cis and trans compartments. Both compartments were filled with PBS electrolyte and electrically connected through a pair of chlorinated silver/silver chloride (Ag/AgCl) electrodes. All measurements were performed inside a Faraday cage at room temperature to minimise external electromagnetic interference. Ionic current recordings were obtained by applying a trans-positive bias voltage of 1.5 V across the nanopore. Current traces were low-pass filtered using a four-pole Bessel filter with a cut-off frequency of 10 kHz and digitised at a sampling frequency of 50 kHz for subsequent analysis. Nanopores with nominal diameters of 60 nm were used for polystyrene bead calibration experiments, whereas 40 nm nanopores were employed for Tau measurements. Data acquisition and event analysis were performed using a custom real-time peak detection program.

### Biophysical Sizing and Shape Modelling of Spherical and Cylindrical Tau

The structural dimensions of Tau assemblies were estimated from their resistive pulse signatures based on the excluded-volume principle of nanopore sensing [41]. Upon translocation through the sensing region, the presence of an analyte displaces electrolyte from the nanopore, leading to a transient increase in pore resistance and a corresponding decrease in ionic current. Under a constant applied potential, the fractional current blockade can be expressed as a function of the resistance increase induced by the translocating particle according to Ohm’s law and classical resistive pulse theory [9]:

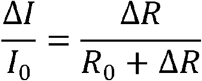

where *I*_0_ represents the open-pore current, Δ*I* denotes the blockade amplitude, *R*_0_ is the open-pore resistance, and Δ*R* corresponds to the additional resistance generated by the excluded electrolyte volume of the analyte.

To relate the measured blockade amplitudes to physical dimensions, compact Tau species were approximated as spherical particles, whereas elongated aggregates were represented using a cylindrical geometry (Fig. 1B). For spherical particles, the particle volume is given by

**Figure 1.**
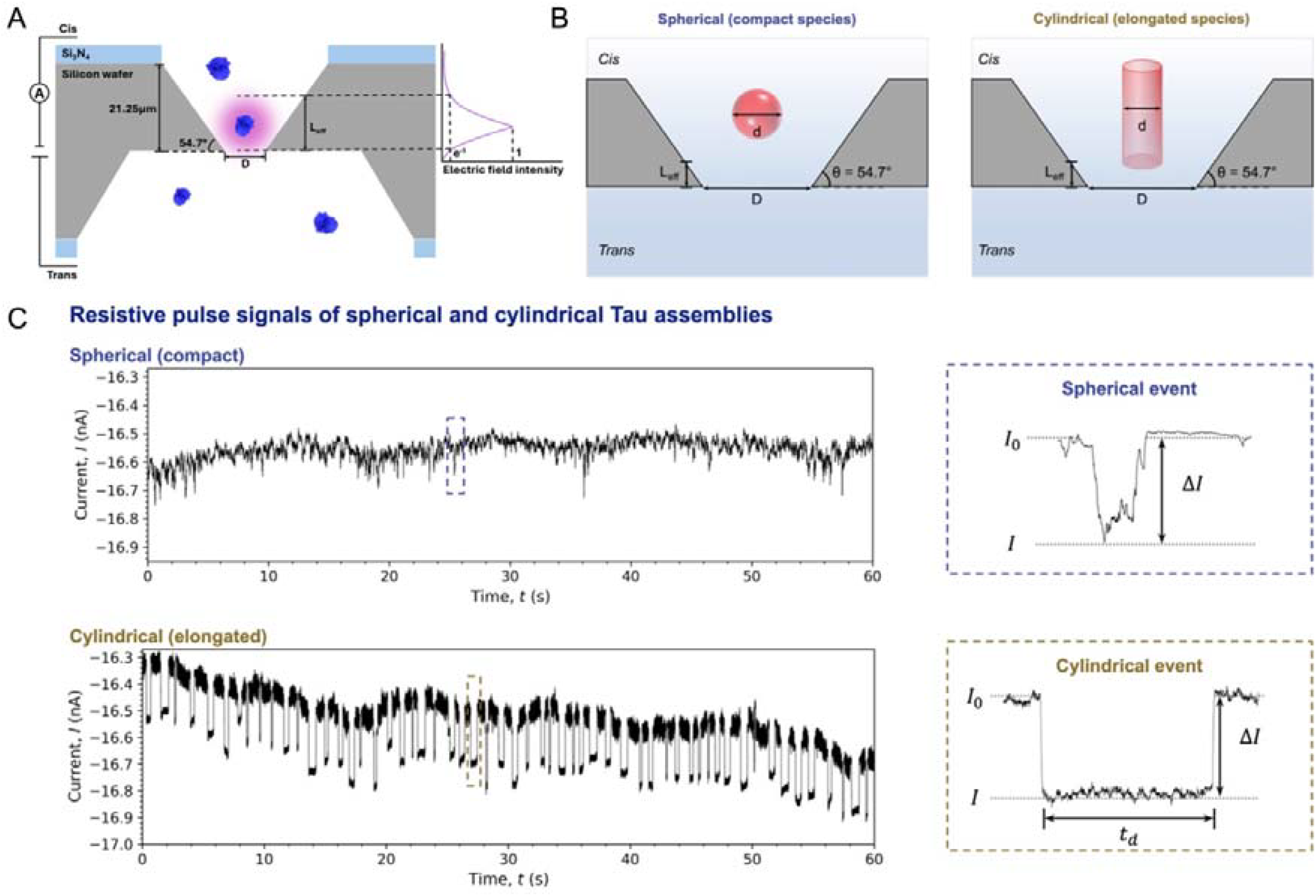
Biophysical sizing framework used for structural characterisation of Tau assemblies. (A) Cross-sectional schematic of the micro-pyramidal silicon nanopore showing the geometrical parameters used in the model. (B) Cross-sectional schematic for compact and elongated Tau species represented by spherical and cylindrical models, respectively. (C) Representative resistive pulse signals of spherical and cylindrical models illustrating the blockade amplitude (*ΔI*) and dwell time (t_d_) extracted from individual translocation events.

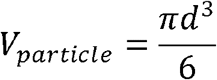

where *d* is the effective particle diameter. Owing to the anisotropic geometry of the micro-pyramidal silicon nanopore, a geometrical correction associated with the 54.7° inclination angle between the silicon crystallographic planes was incorporated according to the pore architecture reported by Yang et al. [26].The effective sensing length was defined as *L*_*eff*_ = *πD*/6, where *D* is the bottom gate diameter of the nanopore.

To improve size estimation for particles occupying a substantial fraction of the pore opening, an analytical shape factor correction *S*(*d*/*D*) inspired by Smythe and Cooke et al. [42, 43] and a spherical boundary-layer correction coefficient were incorporated according to the models proposed by Hurley and Gao et al. [44, 45]:

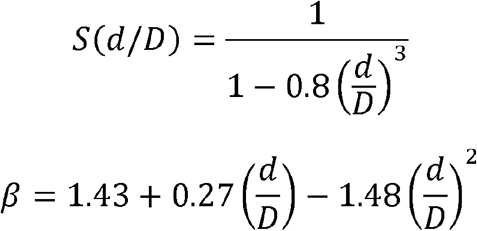

Combining these geometrical correction factors yields the implicit relationship used to estimate the effective diameter of compact Tau species:

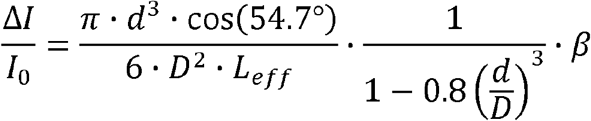

Since the particle diameter appears in multiple terms of the implicit equation, the effective diameter for each event was determined numerically.

For elongated Tau assemblies, the analytes were approximated as rigid cylinders aligned with the pore axis. According to classical resistive pulse theory, the open-pore resistance and blocked-pore resistance can be expressed as [9, 10, 44]

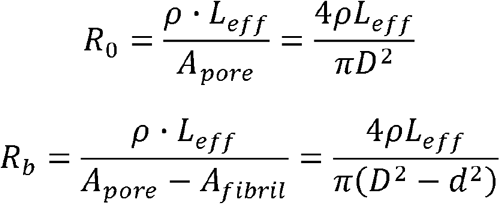

where *ρ* is the electrolyte resistivity, *A*_*pore*_ is the cross-sectional area of the nanopore, and *A*_*fibril*_ represents the projected cross-sectional area of the cylindrical aggregate. Substitution of these resistance terms into the fractional blockade relationship yields:

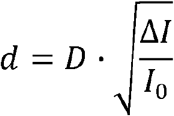

The longitudinal dimension of elongated aggregates was subsequently estimated from the dwell time of each event. Assuming a constant average translocation velocity (*v*) previously calibrated for the micro-pyramidal nanopore platform[26] under fixed experimental conditions, the aggregate length was calculated according to:

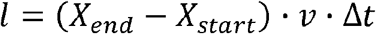

where *X*_*start*_ and *X*_*end*_ correspond to the entry and exit sampling points of the event, *v* is the calibrated translocation velocity, and Δ*t* = 1/*f*, is the sampling interval determined by the acquisition frequency *f*. Under the experimental conditions used in this study ( *v =* 0.005 nm *μ*s^-1^, *f* = 50 kHz), the conversion factor corresponds to 0.1 nm per sampling point, giving:

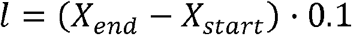

## RESULTS AND DISCUSSION

### Characterisation of Tau Aggregation by TEM

Transmission electron microscopy (TEM) was performed to monitor the morphological evolution of Tau assemblies during heparin-induced aggregation (Fig. 2). Freshly prepared samples collected before heparin addition exhibited largely featureless micrographs with no apparent large aggregates, indicating the absence of pre-existing high-molecular-weight species. After 24 h of incubation, discrete granular structures with heterogeneous morphology became visible, suggesting the formation of intermediate aggregated species. Upon prolonged incubation to 48 h, the samples were dominated by elongated and unbranched fibrillar structures extending over several hundred nanometres in length.

**Figure 2.**
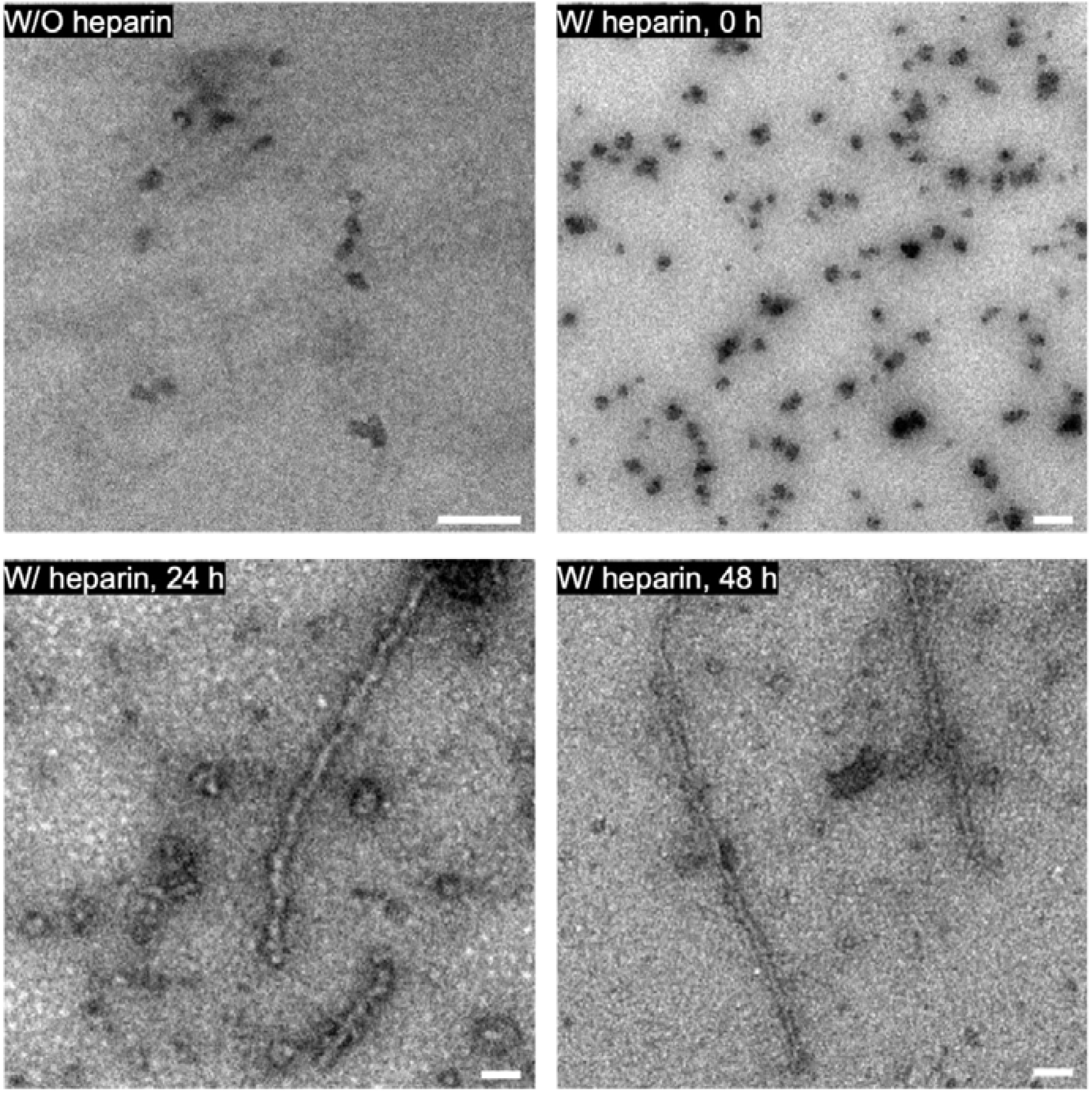
Morphological evolution of Tau assemblies monitored by transmission electron microscopy (TEM). Representative TEM images of Tau samples collected before the addition of heparin and after 0 h, 24 h and 48 h of incubation with heparin. Prior to heparin addition, no apparent aggregates were observed. Immediate incubation with heparin promoted the formation of clustered species, followed by the appearance of protofibrils after 24 h and predominantly mature fibrillar assemblies after 48 h. Scale bars = 100 nm.

The observed morphological progression from dispersed species to granular aggregates and ultimately elongated fibrillar assemblies confirmed the successful induction of Tau aggregation under the experimental conditions employed in this study. These TEM observations provided structural reference information for interpreting the nanopore measurements and supported the use of spherical and cylindrical approximations in the subsequent biophysical sizing analysis. Nevertheless, TEM inherently samples only a limited field of view, restricting the number of particles that can be analysed and limiting the statistical representation of heterogeneous protein populations. While TEM remains indispensable for directly visualising aggregate morphology, the nanopore approach complements this limitation by enabling continuous, label-free analysis of large numbers of individual Tau assemblies in solution, thereby providing a more representative assessment of the heterogeneous Tau population.

### Calibration of the Pyramidal Nanopore Using Polystyrene Beads

Prior to the characterisation of Tau assemblies, the performance of the micro-pyramidal silicon nanopore was evaluated using monodisperse polystyrene beads with a nominal diameter of 50 nm (Yuan Biotech, Shanghai, China). The bead suspension was introduced into a nanopore possessing a bottom gate diameter of 60 nm under an applied bias voltage. Continuous current recordings acquired over a 15 s measurement period exhibited stable baselines and repetitive transient current blockades (drops in ionic current magnitude) corresponding to individual bead translocation events (Fig. 3A). Representative pulse waveforms displayed consistent signal amplitudes and pulse profiles, indicating reproducible particle detection by the nanopore platform (Fig. 3B).

**Figure 3.**
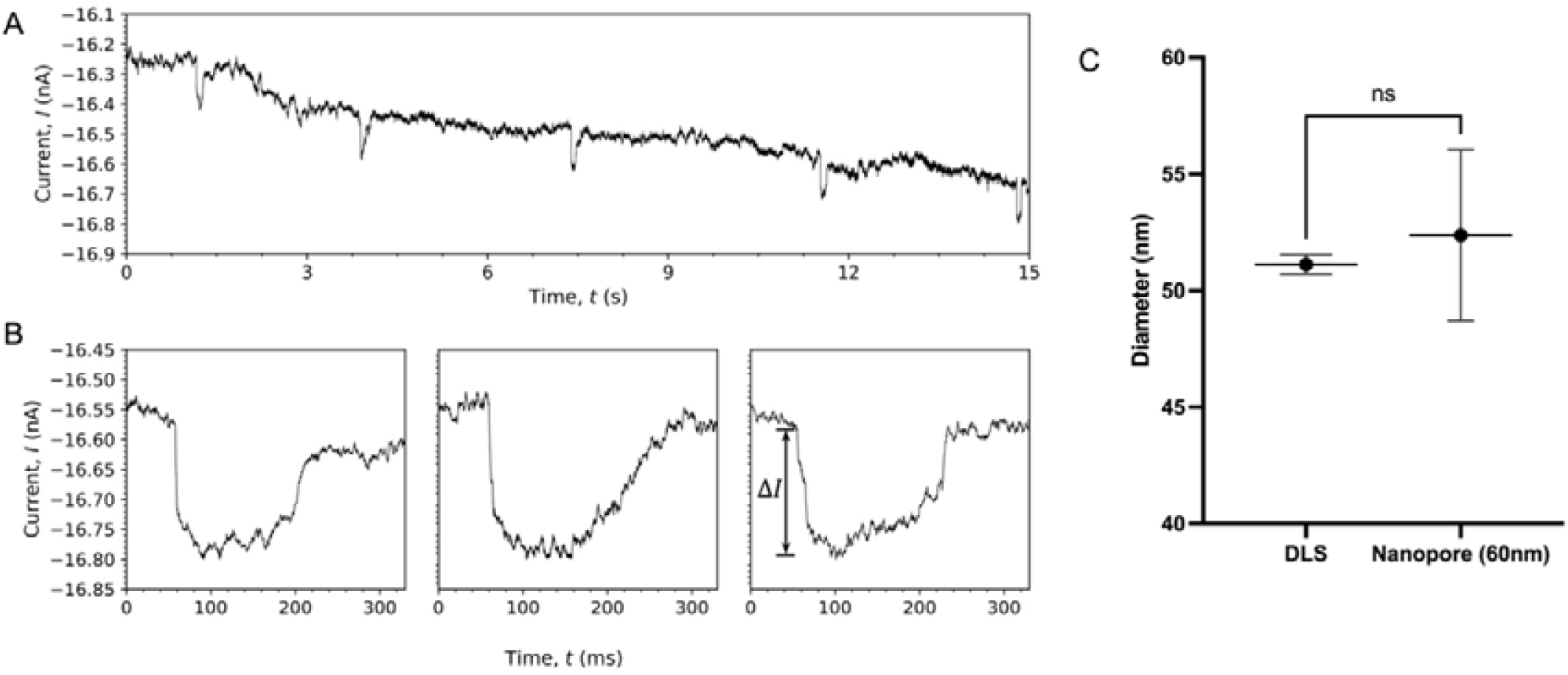
Bead Calibration in Pyramidal Nanopore. (A) A continuous 15-second ionic current recording displaying clean, uniform transient current blockades as the 50 nm polystyrene beads translocate under an applied bias voltage, along with (B) three representative individual resistive pulse profiles. (C) Comparison of diameter distributions of 50 nm beads measured via DLS and the calibrated nanopore method. Statistical analysis using an unpaired t-test revealed no significant difference between the two methods (p = 0.1331, ns).

The measured blockade amplitudes were subsequently analysed using the boundary-layer-corrected pyramidal sizing model described previously to estimate the particle diameter. The calculated bead diameter obtained from nanopore measurements was 52.38 ± 3.66 nm (n = 30), which showed good agreement with the particle diameter, 51.12 ± 0.4106 nm (n = 20), measured by dynamic light scattering (DLS) (Fig. 3C). Statistical analysis using an unpaired t-test revealed no significant difference between the two methods (p = 0.1331). Although the reconstructed bead diameters exhibited a finite distribution, particle diameter was reconstructed indirectly from individual resistive pulse signals using the boundary-layer-corrected geometrical model, such that small variations in the recorded electrical signals contribute to the spread of the reconstructed diameters. The relatively narrow bead distribution establishes the baseline variability of the nanopore sizing approach and provides a useful reference for interpreting the considerably broader distributions observed for Tau assemblies, which primarily reflect the intrinsic heterogeneity of the aggregation process. These results demonstrate that the pyramidal nanopore platform provides reliable size estimation for nanoscale particles and establishes a reference framework for the subsequent characterisation of Tau assemblies. Furthermore, since the underlying geometrical correction factors depend on relative size ratios (*d* / *D*), the bead validation in a 60 nm pore confirms that the mathematical sizing framework scales reliably to the 40 nm pores used for Tau characterisation in the following experiments.

### Nanopore Detection of Tau Assemblies

To investigate the capability of the micro-pyramidal silicon nanopore for monitoring Tau aggregation, ionic current recordings were obtained from freshly prepared Tau samples (with and without heparin) and samples incubated with heparin for 24 hours and 48 hours. Distinct translocation signatures were observed across these conditions, reflecting the structural evolution of Tau assemblies during aggregation.

Freshly prepared Tau samples in the absence of heparin generated frequent downward current blockades with relatively small amplitudes and short dwell times (Fig. 4A), confirming the predominantly monomeric state of the protein. Upon immediate addition of heparin at 0 h (Fig. 4B), infrequent but noticeably sharp, deep downward current spikes appeared. In contrast, samples incubated with heparin for 24 hours produced a marked increase in both event frequency and blockade amplitude (Fig. 4C), indicating the growth of intermediate oligomers and protofibrils. By 48 hours, the samples produced translocation events with noticeably larger blockade amplitudes and prolonged dwell times (Fig. 4D). Individual pulse waveforms from these mature samples displayed broader pulse profiles and extended event durations compared with Tau monomers (Fig. 1C), suggesting the presence of larger, elongated fibrillar assemblies formed during the aggregation process.

**Figure 4.**
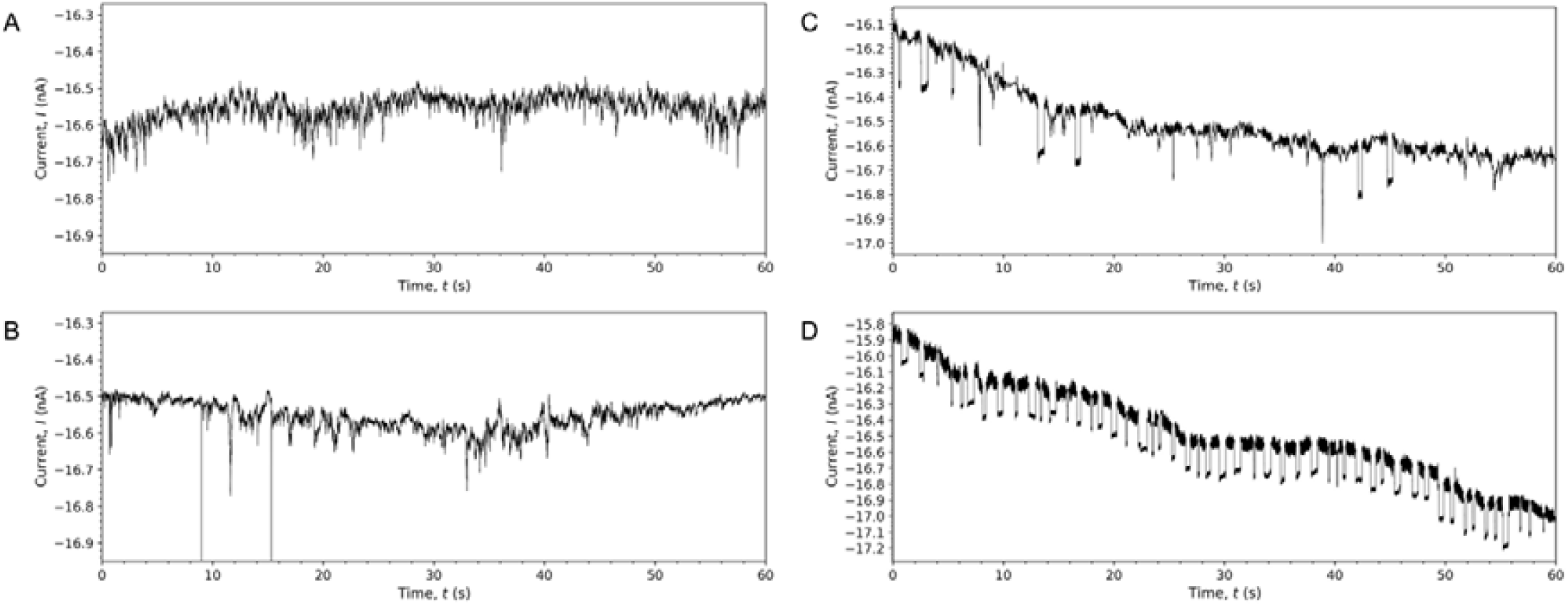
Real-time resistive pulse signatures tracking the time-dependent kinetic progression of Tau aggregation. (A) Continuous 60 s current recording of monomeric Tau in the absence of heparin (t = 0 h). (B) Continuous 60 s recording immediately following heparin addition (t = 0 h), showing sparse, transient nucleation events. (C) Continuous 60 s recording after 24 h of heparin-induced incubation, indicating the emergence of intermediate oligomers and early fibrillar assemblies. (D) Continuous 60 s recording after 48 h of incubation, displaying dense, large-amplitude, and extended-duration blockade pulses characteristic of mature Tau fibrils.

To further compare the electrical signatures obtained from different aggregation stages, the extracted events were analysed according to their fractional current blockades and dwell times. The distributions obtained from heparin-free Tau monomers, intermediate 24-hour assemblies, and 48-hour Tau fibrils samples exhibited distinct characteristics, revealing a clear time-dependent kinetic progression and indicating that the nanopore measurements were sensitive to the structural changes occurring during Tau aggregation. These observations demonstrate that the pyramidal silicon nanopore is capable of detecting aggregation-dependent changes in Tau assemblies through their single-molecule resistive pulse signatures without the need for fluorescent or chemical labels.

### Biophysical Reconstruction of Tau Structural Dimensions

To extract quantitative structural information from the nanopore recordings, the geometrical models described previously were applied to convert the resistive pulse signatures of Tau into the corresponding physical dimensions (Fig. 5). Based on the morphological observations obtained from TEM, freshly prepared Tau species were approximated using the spherical model, whereas elongated assemblies formed after prolonged incubation were analysed using the cylindrical model.

**Figure 5.**
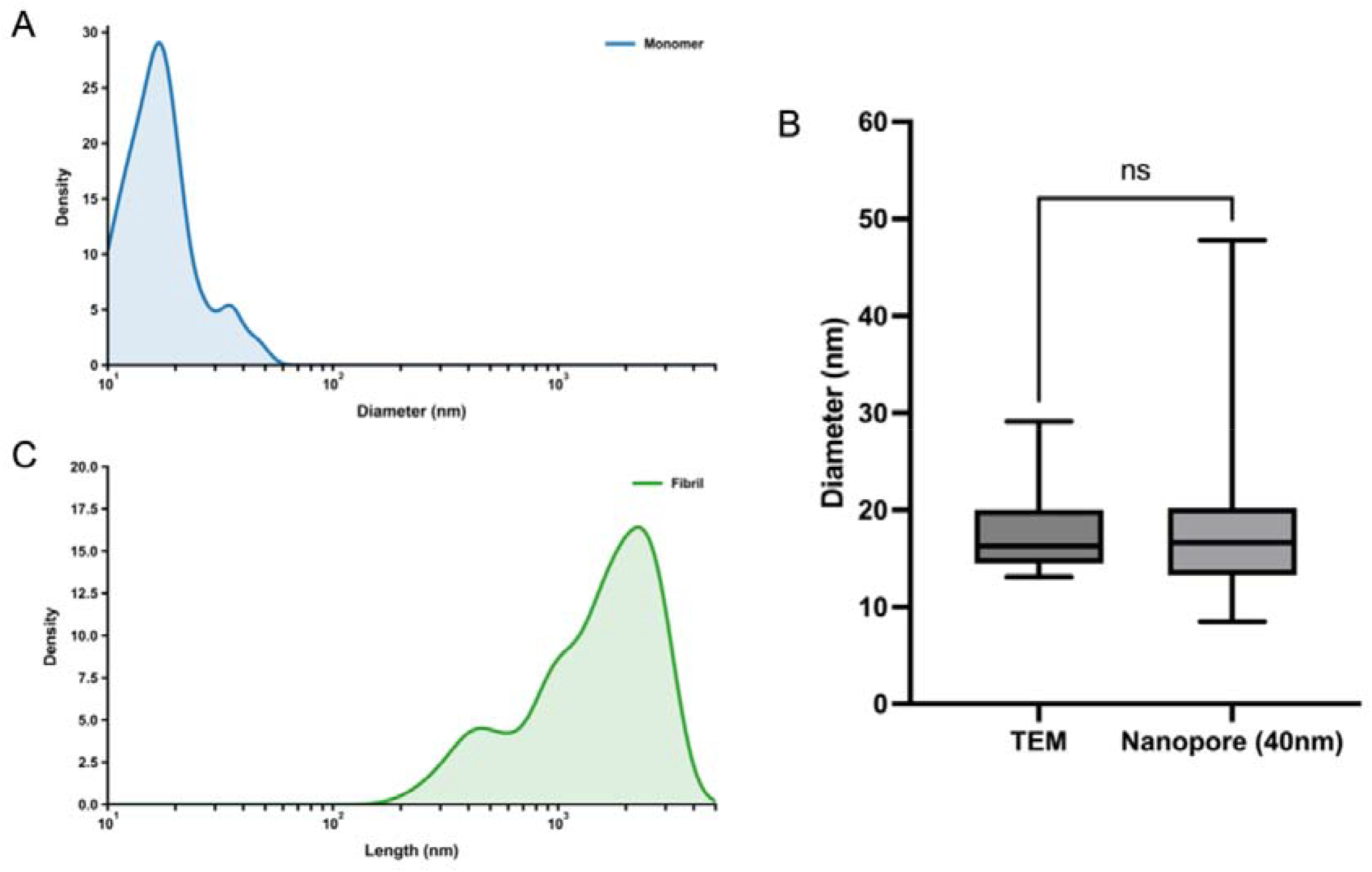
Biophysical reconstruction of Tau structural dimensions from nanopore measurements. (A) Diameter distribution of freshly prepared Tau assemblies reconstructed from fractional current blockades using the boundary-layer-corrected spherical model, revealing compact species with an average diameter of 18.2 ± 7.8 nm (n = 70, N = 3). (B) Comparison of Tau diameters obtained from nanopore measurements and TEM analysis. Statistical analysis using the Mann–Whitney test showed no significant difference between the two methods (p = 0.4237), demonstrating good agreement between electrical measurements and direct structural observations. (C) Length distribution of 48 h Tau assemblies reconstructed using the cylindrical model based on dwell-time analysis. The resulting fibrillar structures exhibited an average length of 1643 ± 842.2 nm (n = 262, N = 3), reflecting the heterogeneous nature of Tau aggregation. Together, these results demonstrate the ability of the pyramidal silicon nanopore platform to estimate structural dimensions of Tau assemblies from single-molecule electrical recordings.

For freshly prepared Tau samples, the measured fractional current blockades were converted into equivalent particle diameters using the boundary-layer-corrected spherical model. The reconstructed diameter distribution exhibited a relatively narrow range centred around 18.2 ± 7.8 nm (n = 70, N = 3) (Fig. 5A), consistent with the predominantly monomeric state of freshly prepared Tau prior to extensive aggregation. To assess the reliability of the nanopore-derived dimensions, the resulting diameter distribution was compared with measurements obtained from TEM. Statistical analysis using the Mann– Whitney test revealed no significant difference between the two methods (p = 0.4237), indicating good agreement between the electrical measurements and direct structural observations (Fig. 5B). Although the reconstructed diameters exhibited a finite distribution, the observed variability likely reflects both the indirect nature of nanopore-based size reconstruction and the inherent conformational heterogeneity of soluble Tau species. The good agreement with TEM measurements indicates that this variability does not compromise the overall reliability of the nanopore-derived structural reconstruction.

For the 48-hour samples, TEM images revealed predominantly elongated fibrillar structures. Therefore, the cylindrical model was employed to estimate the spatial length of individual assemblies from their measured dwell times. The reconstructed length distribution spanned a broad range, with an average length of 1643 ± 842.2 nm (n = 262, N = 3) (Fig. 5C). The broad distribution is consistent with the stochastic nature of heparin-induced Tau aggregation, in which fibrils continuously undergo nucleation, elongation and fragmentation, giving rise to a heterogeneous population of assemblies spanning a wide range of lengths. Consequently, the reconstructed dimensions should be interpreted as representing the evolving distribution of Tau species present at each aggregation stage, rather than the absolute length of individual fibrils. The ability to estimate structural dimensions directly from single-molecule electrical recordings and resolve the evolving distributions of Tau assemblies demonstrates the potential of the pyramidal silicon nanopore platform for label-free characterisation and real-time monitoring of Tau aggregation. When combined with the high spatial resolution of TEM, the nanopore approach provides complementary statistical information from substantially larger numbers of individual translocation events, enabling a more representative assessment of the structural evolution of Tau populations throughout the aggregation process.

### FUTURE PERSPECTIVES

The present study demonstrates that pyramidal silicon nanopores can resolve structural differences in Tau assemblies and reconstruct their dimensions using simplified geometrical models; however, several limitations remain that define clear directions for future development. First, the current calibration strategy relies on 50 nm polystyrene beads measured in a 60 nm pore, whereas Tau measurements were performed in a smaller 40 nm pore to mitigate clogging. This mismatch in calibration and sensing conditions introduces systematic uncertainty in absolute size quantification and highlights the need for size-matched calibration standards, ideally spanning the same dimensional regime as biological analytes.

Future work should focus on expanding the structural resolution capability of the platform. In the present study, spherical and cylindrical approximations were sufficient to describe compact and elongated Tau assemblies; however, intermediate oligomeric or polymorphic species could not be explicitly resolved. Addressing this limitation may require integrating higher-bandwidth signal acquisition, improved signal deconvolution algorithms, or machine learning–based event classification to better capture subtle variations in resistive pulse morphology. In parallel, refinement of nanopore geometry and surface engineering could further reduce event variability and enhance discrimination between closely related aggregation states.

Finally, future developments should aim to bridge the gap between nanoscale sensing platforms and real-time, automated experimental control. This includes the development of closed-loop analytical systems in which AI-assisted data interpretation directly informs measurement strategies, such as adaptive sampling, event-triggered acquisition, or selective target interrogation. Extending such approaches across integrated microfluidic and analytical platforms could ultimately enable more autonomous and high-throughput workflows for biomolecular screening and therapeutic evaluation. As these technologies converge, interdisciplinary collaboration between engineers, biologists, and computational scientists will be essential to translate methodological advances into practical biomedical impact.

## CONCLUSION

In summary, this study demonstrates that pyramidal silicon nanopores can resolve structural heterogeneity in Tau assemblies at the single-molecule level and enable quantitative reconstruction of their dimensions using simplified geometrical models. By combining resistive pulse sensing with a nanoscale confinement architecture, the platform provides a sensitive and label-free approach for distinguishing distinct aggregation states of Tau. To support cross-condition comparability, calibration in this work was performed using nanoparticle standards measured under defined pore geometries, enabling consistent interpretation of resistive pulse signatures across different experimental conditions.

More broadly, this work establishes a foundation for extending nanopore-based sensing toward more complex protein aggregation systems and mechanistic studies of neurodegenerative disease-related species. Future improvements in signal processing, device engineering, and integration with automated analytical frameworks will further enhance structural resolution and throughput, enabling more comprehensive and scalable analysis of protein aggregation dynamics.

## AUTHOR INFORMATION

### Authors

Hoi Lam Cheung - *Ho Sin-Hang Engineering Building, The Chinese University of Hong Kong, Shatin, NT, HK*;

Dehua Hu - *Ho Sin-Hang Engineering Building, The Chinese University of Hong Kong, Shatin, NT, HK*

Jianxin Yang - *Ho Sin-Hang Engineering Building, The Chinese University of Hong Kong, Shatin, NT, HK*

## NOTES

The authors declare no competing financial interest.

## ACKNOWLEDGEMENTS

H.P.H. acknowledges financial support from the Research Grants Council (RGC) General Research Fund (GRF) of Hong Kong (Project No. 14211223). We gratefully acknowledge the Department of Physics at the Chinese University of Hong Kong (CUHK) for providing access to the Transmission Electron Microscopy (TEM) facility.

